# Liquid-Liquid Phase Separation is Driven by Large-Scale Conformational Unwinding and Fluctuations of Intrinsically Disordered Protein Molecules

**DOI:** 10.1101/621714

**Authors:** Anupa Majumdar, Priyanka Dogra, Shiny Maity, Samrat Mukhopadhyay

**Author notes:** Corresponding author: Samrat Mukhopadhyay, Indian Institute of Science Education and Research (IISER) Mohali, Knowledge City, Sector 81, S.A.S. Nagar, Mohali 140306, Punjab, India.

## Abstract

Liquid-liquid phase separation occurs via a multitude of transient, non-covalent, intermolecular interactions resulting in phase transition of intrinsically disordered proteins/regions (IDPs/IDRs) and other biopolymers into mesoscopic, dynamic, non-stoichiometric, supramolecular condensates. IDPs resemble associative polymers possessing stereospecific “stickers” and flexible “spacers” that govern the transient chain-chain interactions and fluidity in phase-separated liquid droplets. However, the fundamental molecular origin of phase separation remains elusive. Here we present a unique case to demonstrate that unusual conformational expansion events coupled with solvation and fluctuations drive phase separation of tau, an IDP associated with Alzheimer’s disease. Using intramolecular excimer emission as a powerful proximity readout, we show the unraveling of polypeptide chains within the protein-rich interior environment that can promote critical interchain contacts. Using highly-sensitive picosecond time-resolved fluorescence depolarization measurements, we directly capture rapid large-amplitude torsional fluctuations in the extended chains that can control the relay of making-and-breaking of noncovalent intermolecular contacts maintaining the internal fluidity. Our observations, together with the existing polymer theories, suggest that such an orchestra of concerted molecular shapeshifting events involving chain expansion, solvation, and fluctuations can provide additional favorable free energies to overcome the entropy of mixing term during phase separation. The interplay of these key molecular parameters can also be of prime importance in modulating the mesoscale material property of liquid-like condensates and their maturation of into pathological gel-like and solid-like aggregates.

## INTRODUCTION

Liquid-liquid phase separation (LLPS) has been recognized as a fundamental phenomenon that drives the assemblage of several proteins and/or nucleic acids within the cells into mesoscopic, dynamic, liquid-like, non-stoichiometric supramolecular assemblies.^1–15^ These biomolecular condensates are non-membrane-bound intracellular bodies, known as membrane-less organelles, that modulate the spatiotemporal localization of different cellular components, organize cellular biochemistry, and regulate a myriad of critical cellular functions. A growing body of intense current research has revealed that intrinsically disordered proteins/regions (IDPs/IDRs) are excellent candidates for LLPS leading to cellular compartmentalization.^1–22^. In vitro, these IDPs recapitulate the behavior of intracellular phase transition and spontaneously demix from a single (miscible) homogenous aqueous phase into a two-phase system comprising a protein-rich dense droplet phase and a light aqueous phase.^18–22^ The thermodynamics of phase separation involves a critical balance between the enthalpy and the entropy of mixing as described by the classic Flory-Huggins theory of polymer phase transition.^23–25^ The enthalpic term associated with the favorable non-covalent, multivalent interchain interactions counteract the entropy of mixing facilitating phase separation of IDPs. These non-covalent interactions include electrostatic, hydrophobic, hydrogen bonding, dipole-dipole, π-π, and cation-π interactions.^24–30^ Therefore, IDPs represent the biological counterparts of associative polymers comprising the intermolecular “stickers” modulating (transient) stereospecific non-covalent interactions and the flexible “spacers” yielding the fluidity that is inherent to a liquid phase.^23,31^ The low-complexity domains often comprising repetitive amino acid sequences offer conformational heterogeneity and flexibility and allow the weak and transient interchain interactions that are crucial for phase separation.^25–32^

While sequence-encoded phase behavior of several IDPs is under intense scrutiny,^25,33–35^ the fundamental molecular mechanism of phase separation remains poorly understood. In this work, we utilize a combination of sensitive readouts of intramolecular proximity and conformational fluctuations to discern the complex interplay of conformational preference, solvation, and dynamics that is critical for driving phase separation. We chose an IDP, namely tau that is predominantly localized in the axons and is associated with Alzheimer’s disease.^36–40^ A highly amyloidogenic 129-residue fragment of human tau consisting of four imperfect 31-residue repeats, namely tau K18, serves as the microtubule binding domain and is critical for aggregation (Figure 1A).^37,38^ Both full-length tau and the tau K18 fragment are known to phase-separate into liquid-like droplets. These droplets are proposed to promote local nucleation of microtubule bundles in neurons (13) and can potentially serve as the precursors of pathological aggregates in Alzheimer’s disease (21, 41).

**Figure 1.**
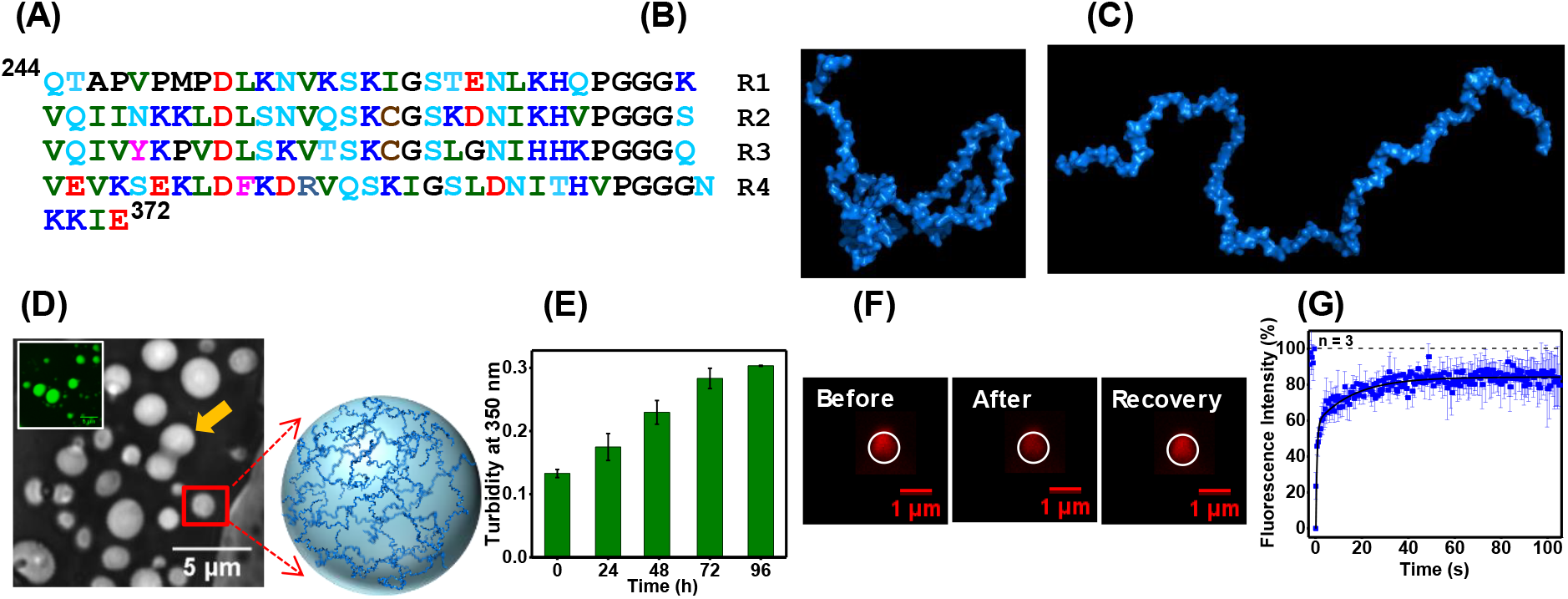
LLPS of tau K18. (A) Amino acid sequence of human tau K18 showing basic (blue), acidic (red), polar (cyan), hydrophobic (olive), cysteine (brown) and all other (black) residues. The NMR structures of (B) a collapsed conformer and (C) an ensemble average conformation^40^ obtained from the Protein Ensemble Database (http://pedb.vib.be) (code: 6AAC) of tau K18 generated using PyMOL (Schrödinger, LLC). (D) An image of droplets with a fusion event shown by an arrow (inset: confocal image of Alexa488-labeled tau K18) and a schematic of a dense protein-rich phase. (E) The changes in turbidity as a function of time (100 μM tau K18 at 37 °C). (F) Images during FRAP measurements of Alexa594-labeled tau K18 droplets. (G) The FRAP kinetics of droplets (mean ± SEM for at least three independent measurements). See Supporting Information for details.

## RESULTS AND DISCUSSION

Tau K18 is an IDP that adopts an ensemble of heterogeneous interconvertible conformers with a preference to collapse in water.^40^ In other words, water serves as a poor solvent for tau K18 as predicted by the PONDR bioinformatics tool^42^ (Supporting Information, Figure S1) and established by NMR (Figure 1B, 1C).^40^ It is known to form liquid droplets at tens of micromolar concentrations under a wide range of solution conditions, with pH varying from 4.8 - 8.8, and at a range of temperature between 25 - 42 °C.^41^ It displays a typical low critical solution transition (LCST) behavior with a characteristic transition temperature ∼15 °C below which it does not phase-separate.^41^ As a prelude, we first confirmed that it forms droplets that recapitulated similar liquid-like characteristics under our experimental condition. Upon incubation at 37 °C, it spontaneously transformed into spherical mesoscopic droplets and the smaller droplets fused to form larger droplets as expected (Figure 1D, 1E). A rapid internal diffusion within a droplet, a key characteristic of a liquid phase, was also confirmed by the fast recovery of fluorescence of Alexa labeled tau K18 in our fluorescence recovery after photobleaching (FRAP) experiments (Figure 1F, 1G).

After establishing that tau K18 forms liquid droplets, we asked the following question: Is there a shift in the conformational distribution during phase separation? In order to answer this, we utilized a sensitive intramolecular proximity probe, namely pyrene that exhibits a strong long-wavelength emission band due to formation of an excimer (excited-state dimer) when two pyrene moieties are placed < 10 Å (43,44). We took advantage of two native Cys residues separated by 31 residues in tau K18 (Figure 1A) and covalently labeled them using thiol-active N-(1-pyrene) maleimide. Then we mixed labeled and unlabeled proteins (1:99) and the mixture formed droplets similar to unlabeled tau K18 (Figure 2A inset). We then monitored the fluorescence characteristics of pyrene as a function of time during phase separation. In the (dispersed) monomeric state, pyrene exhibited a strong excimer band due to highly proximal pyrene moieties in a compact polypeptide chain. This result is consistent with the fact that tau K18 adopts an ensemble of compact structures. During phase separation, the excimer band gradually decreased suggesting that the polypeptide chains adopt more extended structures in the droplet state (Figure 2A). In order to further confirm the decrease in excimer during LLPS, we performed a control experiment by incubating tau K18 at 4 °C which does not promote phase separation due to the LCST behavior^41^. Upon incubation at 4 °C, we did not observe a decrease in the excimer band (Figure 2B). Therefore, our pyrene excimer studies confirmed that phase separation is associated with the conformational expansion of the polypeptide chain from compact globule to extended coil (Figure 2C). It is interesting to note that the interior of a droplet containing a polymer-rich dense phase is known to exhibit a much lower dielectric constant than water^45^ The polypeptide chains can act as a good solvent within the droplets, in which the chains can exhibit extended conformations which allow them to participate in a multitude of non-covalent intermolecular interactions to form physical crosslinks. This structural depiction of the polypeptide chains within the droplets raises two important questions: (i) Does the droplet interior recruit water? (ii) Is there an alteration in the conformational dynamics of the polypeptide chains within the droplets?

**Figure 2.**
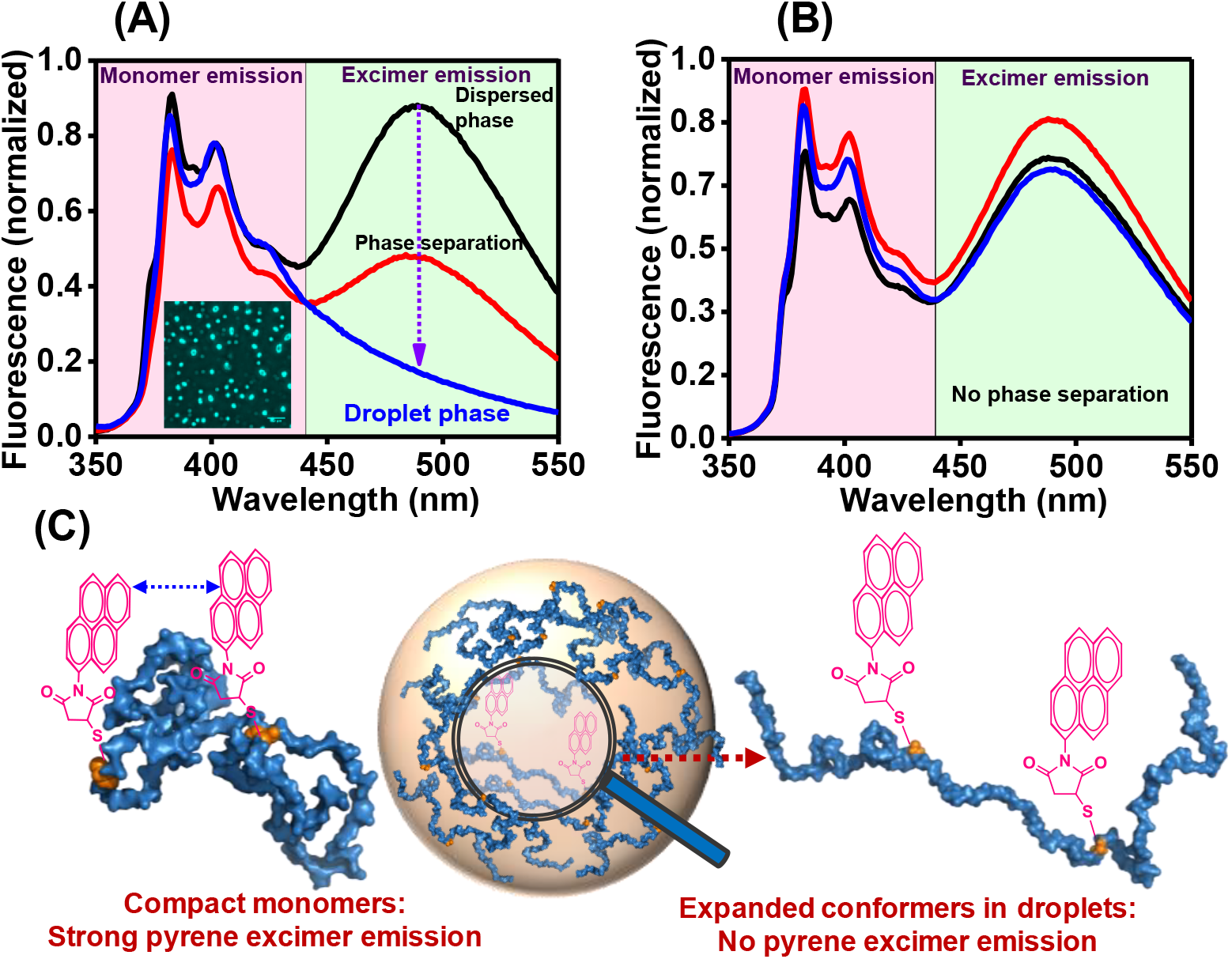
The change in the intramolecular proximity during phase separation. (A) Pyrene emission spectra (1 μM pyrene-labeled + 99 μM unlabeled tau K18) recorded during LLPS at 37°C (black: 0h; red: 24h; blue: 48h after incubation). The inset shows an image of pyrene-labeled tau K18 droplets. (B) Tau K18 incubated at 4°C. The spectra were recorded at 25°C; see Experimental Methods for details. (C) A schematic of the conformational expansion upon phase separation.

We next set out to characterize the inner environment of the droplets. In order to directly probe the solvent environment within the phase-separated droplets, we performed the solvent accessibility measurements using fluorescence quenching experiments of fluorescently-labeled tau K18 with a water-soluble quencher, namely potassium iodide (KI).^43^ For this set of experiments, we covalently labeled Cys with sulfhydryl-reactive fluorescein-5-maleimide. The labeled protein was used as a dopant (0.2%) with unlabeled tau K18. This mixture afforded fluorescent liquid droplets and the presence of a small fraction of labeled protein did not alter the phase separation characteristics (Figure 3A inset). We then estimated fluorescence lifetimes in the absence and in the presence of increasing concentrations of KI to obtain the Stern-Volmer quenching constants and the biomolecular quenching rate constants (Eq. 4 & 5) for monomeric protein, droplets, and free fluorescein as a control (Supporting Information, Table S1). Free fluorescein is close to the upper-limit of high accessibility of water-soluble quencher such as iodide indicated by a steeper slope (Figure 3A). In the monomeric tau K18, fluorescein covalently linked to the protein exhibited much lower water accessibility that is consistent with a more protected environment in the compact ensemble. Surprisingly, we find that water accessibility is much higher in the liquid droplet state compared to the monomeric form of tau K18 (Figure 3A). The order of solvent accessibility is as follows: free dye > droplet > monomer. These results reveal significant chain solvation inside the droplets. This observation is also supported by a considerable red-shift (∼13 nm) observed in the emission spectra of an environment-sensitive label, acrylodan, attached to tau K18 in the droplet state (Supporting Information, Figure S2). Together, these results indicated that tau K18 droplets recruit water molecules and therefore these droplets can qualify as a semi-dilute dense phase. It is interesting to note an intriguing interplay between conformational expansion and water recruitment during phase separation. The solvent quality within the droplets is therefore dictated by the water content as well as the protein-rich milieu giving rise to an environment akin to a polar organic solvent, which is a better solvent than water for the chains to unravel. These extended conformers may promote and modulate transient interchain interactions through rapid conformational fluctuations. We next addressed the dynamical aspects of polypeptide chains within the droplets.

**Figure 3.**
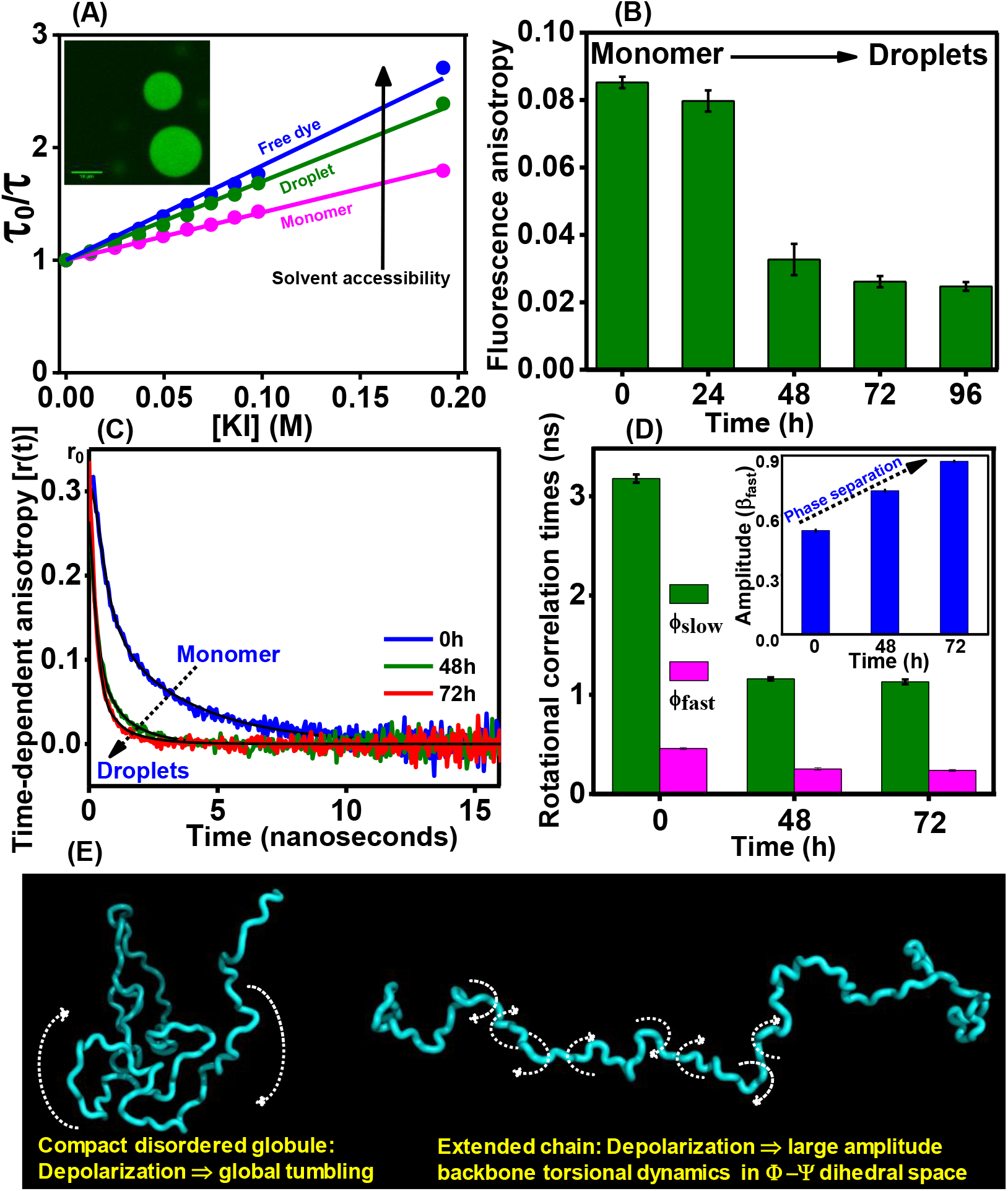
The changes in the solvent accessibility and conformational fluctuations during phase separation. (A) The Stern-Volmer plots of free fluorescein (blue) and monomeric (pink) and droplets formed after 72h (green) from fluorescein-labeled (0.2%) tau K18 by plotting the ratio of mean fluorescence lifetimes in absence and presence of KI (Eq. 4). The inset shows a confocal image. (B) Steady-state fluorescence anisotropy of fluorescein-labeled tau K18 during LLPS. (C) Time-resolved fluorescence anisotropy decays (blue: 0h; green: 48h and red: 72h) The black lines are fits using the biexponential decay of depolarization kinetics (Eq. 8). See Methods for data analysis and Table S2 for recovered parameters associated with depolarization kinetics. (D) Changes in the rotational correlation times (inset: changes in the amplitude of fast correlation time) during phase transition (mean ± SEM for three independent measurements). (E) The molecular origins of picosecond fluorescence depolarization kinetics of collapsed and extended IDPs.

In order to study the conformational dynamics, we measured fluorescence polarization (anisotropy) of fluorescein-labeled tau K18. Fluorescence anisotropy is related to the rotational flexibility and increases during formation of supramolecular assemblies due to the restricted mobility within the assemblies.^43,46^ Therefore, a transition into a condensed phase is expected to dampen the mobility and should increase the anisotropy. We first carried out the steady-state fluorescence anisotropy measurements. Surprisingly, we observed a sharp decrease in the anisotropy value from the monomeric form to the droplet state (Figure 3B). In order to rule out any turbidity-induced artifact, we also measured anisotropy at various dilutions. The measured anisotropy remained unaltered upon dilution indicating that the observed drop in the anisotropy is associated with an increase in the rotational flexibility within the droplets (Supporting Information, Figure S3). Additionally, incubation of tau K18 at 4 °C that inhibits phase separation did not exhibit any drop in the anisotropy (Supporting Information, Figure S4). We further corroborated this observation by using a different label, acrylodan, which showed a similar decrease in the anisotropy upon phase transition (Supporting Information, Figures S5). Taken together, this set of experiments revealed that phase separation is associated with a decrease in the anisotropy. This unusual observation might indicate a more rapid large-scale reorientation of the polypeptide chains within the droplets.

The steady-state fluorescence anisotropy values report average rotational flexibility and do not distinguish between the different dynamical molecular events within a polypeptide chain. On the contrary, the highly-sensitive picosecond time-resolved fluorescence anisotropy measurements allow us to discern the dynamical events that are responsible for fluorescence depolarization from the (fundamental) time-zero anisotropy (r_0_).43,46 The depolarization kinetics can typically be described by a biexponential decay function comprising a (fast) sub-nanosecond rotational correlation time (ϕ_fast_) and a slow rotational correlation time (ϕ_slow_) (Figure 3C and Eq. 8). The ϕ_fast_ represents the local rotational mobility of the attached fluorophore, whereas the molecular origin of the ϕ_slow_ depends on the conformational characteristics of macromolecules. Our previous studies have shown that for compact IDPs, the ϕ_slow_ corresponds to the global tumbling of the collapsed globules depending on the hydrodynamic volume as governed by the Stokes-Einstein relationship.^44^ On the contrary, for the expanded IDPs, the ϕ_slow_ represents a size-independent characteristic timescale (∼1.3 ns) that quantifies the depolarization component arising due to a collective segmental torsional mobility of the Ramachandran Φ-Ψ dihedral angles of the extended polypeptide chain (Figure 3E).^46,47^ Tau K18 exhibited a typical biexponential decay both in the monomeric and in the droplet states (Figure 3C). In the monomeric form, the ϕ_slow_ was ∼ 3 ns that corresponds to the rotational diffusion of a partially collapsed state of tau K18 that is consistent with our excimer data and previous studies.^40^ Upon phase transition, the ϕ_slow_ became much shorter (∼ 1.1 ns) as shown by more rapid depolarization kinetics (Figure 3C, 3D). This relaxation time represents the backbone torsional mobility in the dihedral angle space of expanded disordered conformations.^47^ Additionally, the amplitude associated with the fast correlation time corresponding to the local dynamics also increased during phase separation (Figure 3D inset). These results showed that the extended polypeptide chains within the droplets undergo extensive dihedral rotations on a timescale that is close to the characteristic backbone torsional relaxation expected for expanded chains in good solvents (Figure 3E). Therefore, our finding provides a compelling evidence for rapid large-amplitude torsional angle fluctuations within the liquid droplets.

## CONCLUSION

In conclusion, our study directly captures an intriguing symphony of key molecular shapeshifting events associated with significant chain expansion, solvation, and fluctuations during phase separation (Figure 4). Our pyrene excimer studies allowed us to directly observe the conformational expansion during phase transition. This expansion is possibly governed by a better solvent quality within the droplets compared to the bulk aqueous phase. These extended conformers can be formed via conformational selection and/or transition and can allow the transient intermolecular contacts through a multitude of weak interactions forming a network of physical crosslinks. Tau K18 exhibits an LCST phase transition that can originate from hydrophobic interactions responsible for the entropic release of water molecules.^25,41,48^ The sequence has several hydrophobic residues that can potentially act as the “stickers” in phase separation mediated by entropic gain. Such a mechanism has been proposed for an entropy-driven coacervation that retains the intrinsic disorder in the phase-separated droplets.^49^ Our results also show that the chain expansion can allow the polar and charged residues to get solvated as demonstrated by higher solvent accessibility in droplets compared to monomers. This favorable chain solvation, presumably in the vicinity of the flexible linkers, leads to the recruitment of water in the droplet resembling a semi-dilute polymer phase. Therefore, water might play an intriguing dual role; entropic release can promote hydrophobically-driven phase transition and the polar regions of the protein get highly solvated that is essential for the spacers to undergo rapid chain fluctuations maintaining the internal fluidity that is essential for a liquid-like interior of the droplets. The abundance of glycines (PGGG motifs) in the form of the spacers in the sequence might favor both conformational expansion and rapid torsional fluctuations on a characteristic timescale within the droplets.^30^ We propose that chain unwinding and fluctuations can increase the chain entropy within the droplets. Additionally, these large-scale conformational fluctuations can also promote interchain interactions that can modify the enthalpic contribution to counter the entropy of mixing term.^19^ A modification of the Flory-Huggins theory in the semi-dilute concentration regime was performed by incorporating a three-body term and density fluctuations.^50,19^ Molecular simulations suggested that such large-scale chain fluctuations can generate a larger pervaded overlap volume that facilitates favorable intermolecular interactions.^19^ The linker flexibility can be a critical regulator of weak multivalent interactions in the liquid phase and can govern the material-state of biomolecular condensates.^51^ Our picosecond time-resolved fluorescence depolarization measurements allowed us to directly capture the unique dynamic signatures of the polypeptide chains within the phase-separated droplets. Rapid conformational fluctuations can provide a temporal control of the making and the breaking of stereospecific, intermolecular, noncovalent contacts between the “stickers” on a characteristic timescale that can further allow remaking of the new contacts with different polypeptide chain partners within the droplets. These temporal fluctuations resulting in the relay of making-and-breaking of intermolecular contacts can be crucial for maintaining the liquid-like property in the dynamic assembly. Taken together, results from our studies illuminate the key molecular drivers of phase separation that can be of great importance in the context of functional aspects of a broad range of IDPs involved in cell physiology. Additionally, the interplay of chain expansion, solvation, and fluctuations can serve as the critical molecular parameters to modulate the mesoscale material property of liquid droplets and their maturation into viscoelastic gel-like and pathological solid-like aggregates found in deadly neurodegenerative diseases.

**Figure 4.**
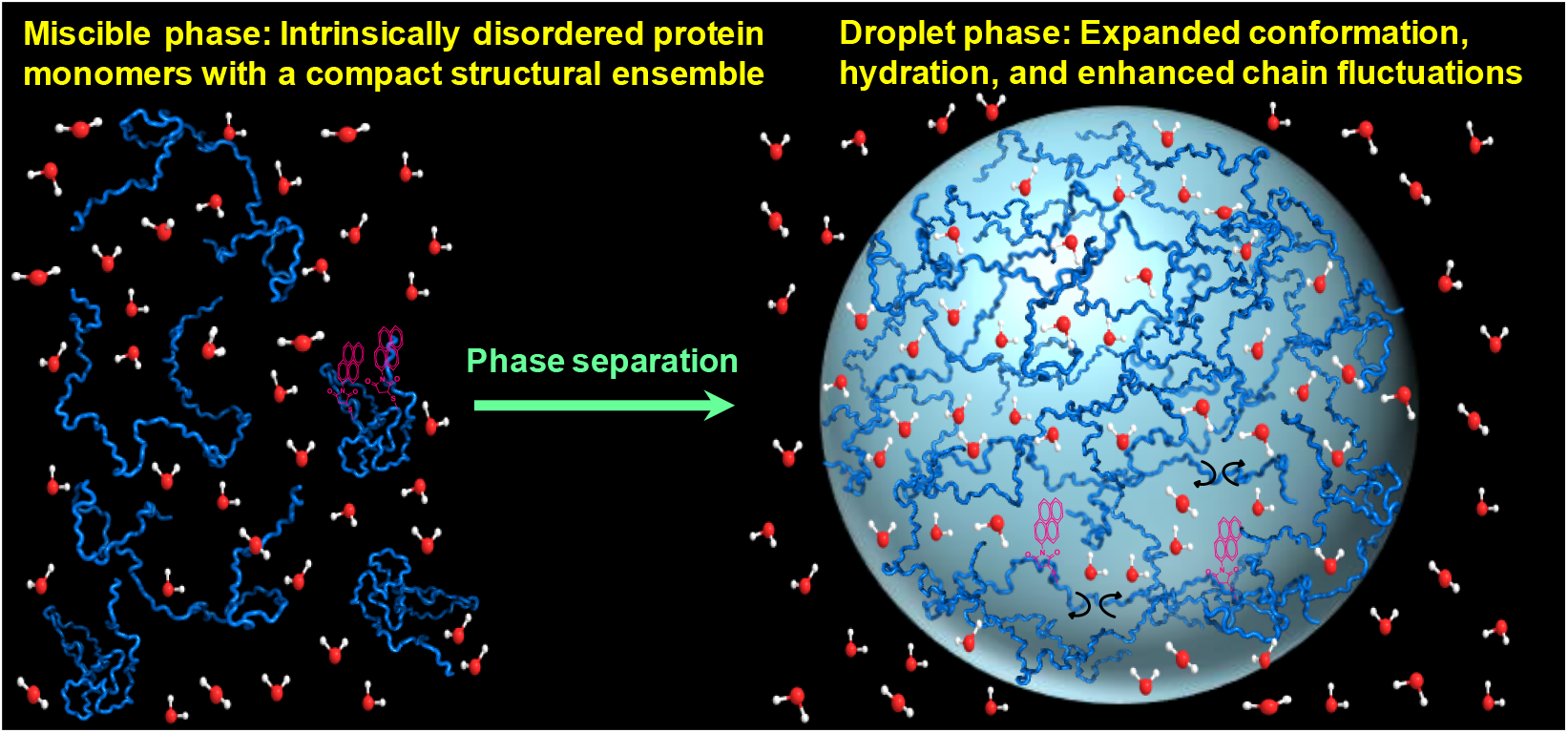
A depiction of molecular shapeshifting events involving chain expansion, hydration, and fluctuation upon liquid-liquid phase separation. The conformational unwinding is recorded by the decrease in the intramolecular proximity using excimer emission. The solvent accessibility studies showed chain solvation and water recruitment in the droplets. The picosecond fluorescence depolarization studies revealed the increase in the conformational fluctuations within the liquid droplets.

## MATERIALS AND METHODS

### Sample preparation

Tau K18 was expressed in *Escherichia coli* and purified using our reported protocol.^52^ The liquid droplets were formed using the procedure described previously.^41^ The experimental details of expression, purification, fluorescence labeling, droplet formation, turbidity assay, confocal microscopy, and FRAP are mentioned in Supporting Information text.

### Steady-state fluorescence measurements

All steady-state fluorescence experiments were performed on a Fluoromax-4 spectrofluorometer (Horiba Jobin Yvon, NJ). All the measurements were made at 25 °C. The pyrene- and fluorescein-labeled tau K18 samples were monitored using excitation wavelengths of 340 nm and 485 nm, respectively. For acrylodan-labeled samples, the excitation wavelength was 375 nm. The steady-state fluorescence anisotropy (*r*_ss_) data were recorded at the respective emission maxima and is given by:

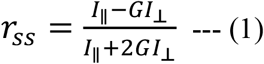

where *I*_||_ and *I*_⊥_ are the recorded fluorescence intensities when the emission polarizer is oriented parallel and perpendicular to the excitation polarizer, respectively, and *I*_⊥_ was corrected using the corresponding G-factor.

### Time-resolved fluorescence measurements

The time-resolved fluorescence measurements were made using a time-correlated single photon counting (TCSPC) setup (Fluorocube, Horiba Jobin Yvon, NJ). All the data were acquired at 25 °C. The samples were excited using a 485-nm NanoLED picosecond diode laser. The instrument response function was recorded using a diluted colloidal suspension of silica and the full-width at half-maxima was estimated to be ∼ 270 ps. The emission monochromator was set at the emission maxima using a fixed bandpass of 8 nm. For fluorescence lifetime measurements, the emission polarizer was set at the magic angle (54.7º) orientation with respect to the excitation polarization. The fluorescence intensity decays [*I*(*t*)] could be fitted to a typical biexponential decay kinetics as follows:

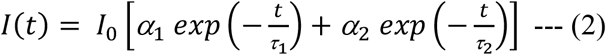

where *I*_0_ is the time-zero intensity, *α*_1_ and *α*_2_ are the amplitudes associated with the lifetime components, *τ*_1_ and *τ*_2_. The amplitude-weighted average mean fluorescence lifetimes were estimated using the following relationship:

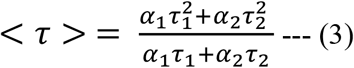

### Estimation of solvent accessibility using fluorescence quenching measurements

The solvent accessibility was measured by time-resolved fluorescence quenching experiments using fluorescein-labeled tau K18 and a potent water-soluble quencher, potassium iodide (KI). Droplets were formed using 0.2% fluorescein-labeled tau K18 (200 nM) mixed with a large excess (99.8 μM) of unlabeled tau K18. The mean fluorescence lifetimes were measured in the presence of increasing concentrations of KI. The time-resolved fluorescence data were recorded analyzed as described above. The ratio of the mean fluorescence lifetimes in the absence and in the presence of quencher [<*τ*_0_>/<*τ*>)], both in the monomeric and in the protein-rich droplet phase, were plotted against the concentration of KI ([*Q*]). The Stern-Volmer quenching constants recovered using the following relationship (Equation 4). The bimolecular quenching rate constant, a quantitative measure of the solvent accessibility, was estimated using Equation 5.

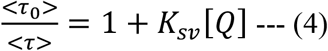

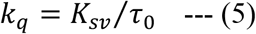

### Fluorescence depolarization kinetics using picosecond time-resolved fluorescence anisotropy measurements

The time-resolved fluorescence anisotropy decay measurements of the samples were performed using a TCSPC setup (Fluorocube; Horiba Jobin-Yvon, NJ). The samples were excited using a 485-nm picosecond diode laser and the emission monochromator was fixed at the corresponding fluorescence emission maxima. The parallel [*I*_||_(*t*)] and perpendicular polarized [*I*_⊥_(*t*)] intensity decays were collected by toggling the emission polarizer between 0° and 90° orientations with respect to the excitation polarization. The *I*_⊥_(*t*) was corrected using a G-factor that was estimated using a solution of the free fluorescein in water. The *I*_||_(*t*) and *I*_⊥_(*t*) were analyzed globally and fitted using the following relationships:^46,47^

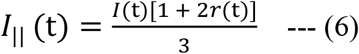

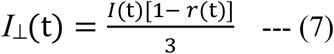

where *r*(*t*) represents the time-resolved anisotropy decay from the (fundamental) time-zero anisotropy (*r*_0_). The anisotropy decay profiles exhibited a typical biexponential depolarization kinetics (Equation 8) giving rise to two rotational correlation times, *ϕ*_fast_ and *ϕ*_slow_ and two corresponding amplitudes *β*_fast_ and *β*_slow_. The *ϕ*_fast_ represents the local rotational dynamics of the probe, whereas, the *ϕ*_slow_ represents the global/segmental dynamics of the protein.^46,47^

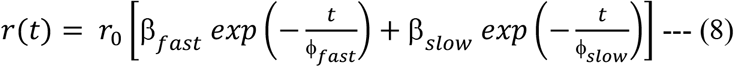

## SUPPORTING INFORMATION

Details of experimental procedures, Supporting Figures S1-S5, and Tables S1-S2 (PDF)

## ACKNOWLEDGEMENTS

We thank IISER Mohali, Department of Science and Technology (SERB National Postdoc Fellowship to A.M.; INSPIRE Fellowship to P.D.; NanoMission grant to S.M.), and Ministry of Human Resource Development (Centre of Excellence grant to S.M.), Govt. of India, for financial support, Prof. E. Rhoades (University of Pennsylvania) for donating tau K18 plasmid, Dr. M. Sharma, Mr. D. Dwivedi, and Ms. R. Marwaha (IISER Mohali) for confocal microscopy, Prof. N. Periasamy (Retd. TIFR Mumbai) for providing the fluorescence decay analysis software, and Dr. M. Bhattacharya (Thapar Institute) and the members of the Mukhopadhyay lab for their valuable comments.

The authors declare no conflict of interest.

