## Supplementary Information for "Liquid-Liquid Phase Separation is Driven by Large-Scale Conformational Unwinding and Fluctuations of Intrinsically Disordered Protein Molecules"

##### Materials

All of the chemicals used for preparing buffer solutions, such as, sodium phosphate monobasic dihydrate, Tris (2-carboxyethyl) phosphine hydrochloride (TCEP), MES (2-(N-Morpholino)ethanesulfonic acid hydrate, EDTA (Ethylenediaminetetraacetic acid), Magnesium chloride hexahydrate, DTT (DL-Dithiothreitol) were of highest purity grade obtained from Sigma Aldrich (St. Louis, MO). The fluorescent probes, namely, fluorescein-5-maleimide, N-(1-pyrene) maleimide, Acrylodan (6-Acryloyl-2-Dimethylaminonaphthalene), AlexaFluor 488 C5-maleimide, and AlexaFluor 594 C5-maleimide were purchased from Molecular Probes, Invitrogen. The free fluorescein dye was purchased from Fluka Analytical. SP Sepharose resin used for protein purification and PD-10 columns were purchased from GE Healthcare Life Sciences (USA). The protein concentrators and filters were procured from Merck Millipore. A Metrohm 827 lab pH meter was used to adjust the final pH (± 0.01) of all the buffer solutions prepared in Milli-Q water and filtered before use.

##### Expression and Purification of tau K18

Tau K18 was expressed in *Escherichia coli* BL21(DE3) and purified using the procedure described previously (51). Briefly, using a lysis buffer of pH 8, (50 mM Tris, 150 mM NaCl, 10 mM EDTA), the cells were lysed by boiling it for half an hour at a 100 ˚C. The lysate was then centrifuged at 11,500 rpm at 4 ˚C for 30 min, following which the supernatant was treated with 136 µL/mL of 10% streptomycin sulfate and 228 µL/mL of glacial acetic acid for the precipitation of DNA. After the removal of DNA by further centrifugation at 11,500 rpm at 4 ˚C for 30 min, the supernatant collected was mixed with equal volumes of saturated ammonium sulfate solution (4 M) and kept for 2-3 h at 4˚C for salting out the protein. The protein pellet obtained upon centrifugation was washed with 100 mM ammonium acetate and 100% chilled ethanol and kept for drying at 37 ˚C overnight. After the complete removal of any traces of ethanol, the dried pellet was dissolved in the native buffer [20 mM MES, 1 mM EDTA, 2 mM DTT and 1 mM $MgCl_2$, (pH 6.8)] and purified on a cation exchange SP Sepharose column (GE Healthcare Life Sciences). The fractions collected were further dialyzed overnight in the native buffer [20 mM MES, 1 mM EDTA, 2 mM DTT and 1 mM $MgCl_2$ (pH 6.8)] to remove the excess salt, following which the protein was stored at -80 ˚C. The purity of the protein was analyzed by SDS-PAGE.

**Fluorescence labeling of tau K18 with *N*-(1-pyrene) maleimide**

The two native cysteine residues in tau K18 were covalently labeled with a fluorescent probe, *N*-(1-pyrene) maleimide, under denaturing conditions. Initially, tau K18 was incubated in a 6 M GdmCl (50 mM phosphate, 1 mM TCEP) buffer of pH 7 and kept overnight at 4 ˚C to ensure its complete denaturation. From a freshly prepared stock (50 mM) of pyrene-maleimide (in DMSO), around 5 μL of the dye was added after every 10 min to the reaction mixture to avoid precipitation. The labeling was carried out under constant stirring of 1500 rpm at 37 ˚C for 4 hours, maintaining a final ratio of 1:10:30 for tau K18: TCEP: dye. At the end of the reaction, tau K18 was eluted with 50 mM sodium phosphate (0.5 mM TCEP) buffer of pH 8.8 using a PD-10 column to remove the unreacted dye. The initial fractions of the eluted protein were pooled together, and the concentration of the labeled protein was calculated using a molar extinction coefficient of 40,000 $M^{-1}cm^{-1}$ at 340 nm. The labeled protein was used in doping concentrations (1%) for the experiments (labeled: unlabeled 1: 99), so as to ensure that the observed excimer fluorescence was due to intramolecular interaction as opposed to intermolecular interaction.

**Fluorescence labeling of tau K18 with other thiol-active fluorescent dyes**

Tau K18 was covalently labeled, non-selectively, at the two cysteine residue positions under native conditions (20 mM MES, 2 mM DTT, 1 mM $MgCl_2$, 1 mM EDTA, pH 6.8) with fluorescein-5-maleimide (F-5-M) (0.5 equivalents), AlexaFluor 488 C5-maleimide (2 equivalents), AlexaFluor 594 C5-maleimide (2 equivalents), Acrylodan (2 equivalents), respectively. The reaction mixtures were kept at room temperature under constant stirring of

6 rpm for 2 hrs. After the completion of the reactions, PD-10 columns were used to remove the free dye and the labeled proteins were eluted using 50 mM sodium phosphate (0.5 mM TCEP) buffer at pH 8.8. The concentrations of the labeled proteins mixed with the unlabeled reactions for setting up droplet reactions have been mentioned in the respective figure legends.

**Liquid Droplet Formation**

For all the measurements, tau K18 droplets were formed upon incubation of 100 µM tau K18 (50 mM sodium phosphate, pH 8.8, 0.5 mM TCEP at 37 ˚C) as described previously (41). The samples were prepared in autoclaved 1.5 mL micro-centrifuge tubes where the buffer solutions were carefully added to the protein and care was taken to avoid the formation of any bubbles in the solution. Phase separation of tau K18 was also monitored at pH 7.4 and 6.8 that resulted in droplet formation. No droplet formation was observed at 4 ˚C that was expected because of the low critical solution transition behavior of tau K18 (40). For all spectroscopic experiments, droplet formation was always confirmed by performing turbidity assays and confocal microscopy.

**Turbidity assay**

The turbidity measurements were performed by recording the optical density of the protein solutions (~ 150 μL) at 350 nm using a 96-well optical bottom NUNC plate on a Thermo Scientific Multiskan Go plate-reader instrument. All the data were recorded in triplicates.

**Confocal Microscopy**

Images of phase-separated liquid droplets of tau K18 were imaged on the Olympus FLUOVIEW confocal laser scanning microscope (Model no. FV10i) using a 60x oil-immersion objective (Numerical aperture: 1.35). The tau K18 droplet reactions were divided into ~ 10 μL aliquots for visualization at different time-points of the reaction. From each of these aliquots, around 5-6 μL was transferred onto a fresh microscope slide (Fisher Scientific 3″ x 1″ x 1 mm) which was then immediately covered with a circular coverslip and sealed with nail-paint at either of the two opposite edges. This was done to avoid evaporation of the solution during image acquisition. The labeled tau K18 protein was mixed with unlabeled proteins at specific ratios as mentioned in the main manuscript. The excitation sources used for visualizing the pyrene-1-maleimide, fluorescein-5-maleimide/AlexaFluor 488 C5-maleimide, and the AlexaFluor 594 C5-maleimide labeled droplets were the 405 nm (17.1

mW), 473 nm (11.9 mW), and 559 nm (15 mW) laser diodes, respectively, and the corresponding emission wavelengths were 461, 520, and 618 nm, respectively. The images were then imported and analyzed in the ImageJ software. (NIH, Bethesda, MD, USA).

**Fluorescence Recovery after Photobleaching (FRAP) Measurements**

The mobility of AlexaFluor 594 C5-maleimide labeled protein molecules within the liquid droplets was assessed using FRAP measurements, carried out on a Confocal Laser Scanning ZEISS 710 Microscope having a Plan Apochromat 63x oil-immersion objective (Numerical aperture 1.4) along with a high-resolution monochrome cooled AxioCamMRm Rev. 3 FireWire(D) camera. The ZEN Pro 2011(ZEISS) software was used for image acquisition as well as for recording the FRAP profiles. For recording the FRAP profiles of the droplets, a rectangular spot (area ~ 0.05 – 0.1 $\mu m^2$) was bleached using a 561 nm laser (20 mW laser power) for ~ 0.5 – 1 s and the fluorescence recovery time-lapse profiles were acquired at a 0.5 s frame rate with a total of 400 frames. The fluorescence intensity recovery profiles were plotted as a function of time using the OriginPro 8.5.1 plotting and analysis software.

### Supporting Information Figures & Tables

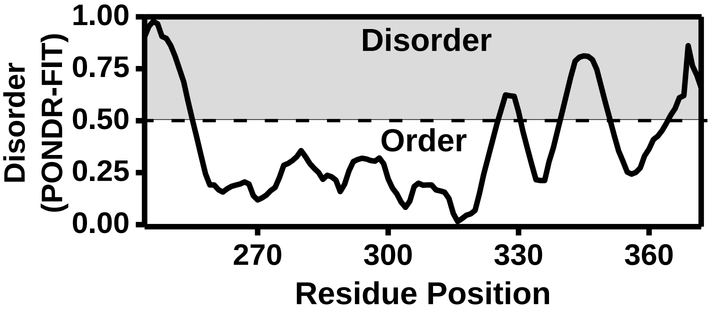


**Figure S1.** PONDR plot predicting disorder in the tau K18 sequence generated using (http://www.pondr.com/) and plotted using OriginPro 8.5.1 software.

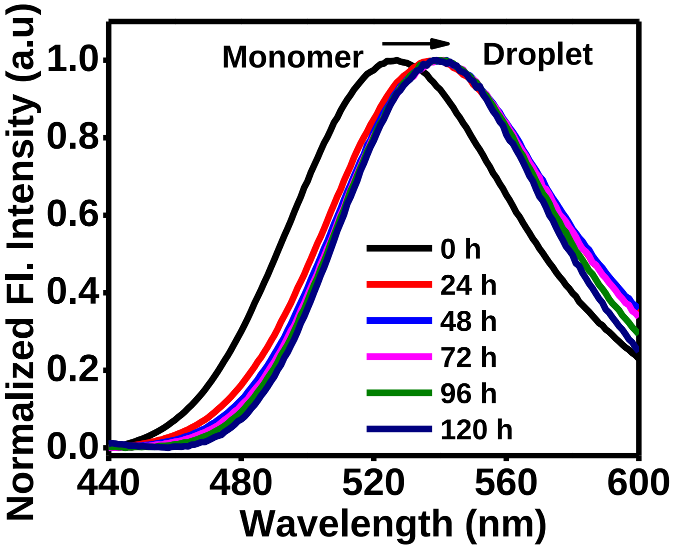


**Figure S2.** Fluorescence emission spectra of solvent-sensitive acrylodan labeled tau K18 (100 μM; ~ 50 % labeling efficiency), at the two native cysteines (positions 291 and 322), showing a significant red-shift as a function of increased droplet formation due to LLPS at 37 °C.

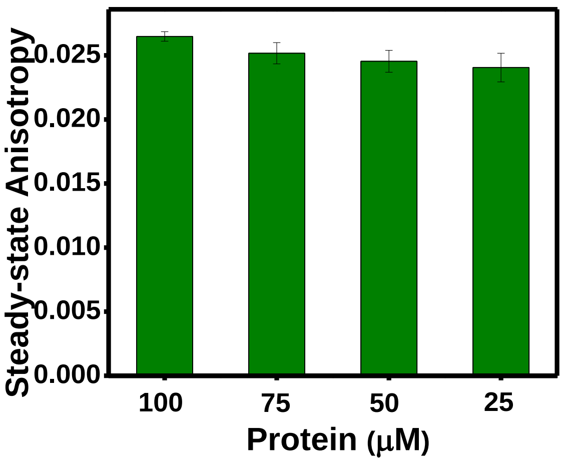


**Figure S3.** Steady-state fluorescence anisotropy of fluorescein-labeled tau K18 recorded as a function of serial dilution of the droplets formed from 100 μM protein (0.2 % labeled) after 72 h.

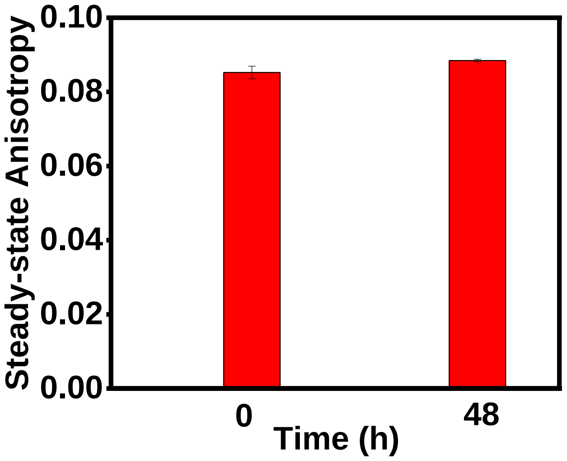


**Figure S4**. No decrease in the fluorescence anisotropy of fluorescein-labeled tau K18 was observed after incubating at 4 °C that does not result in droplet formation.

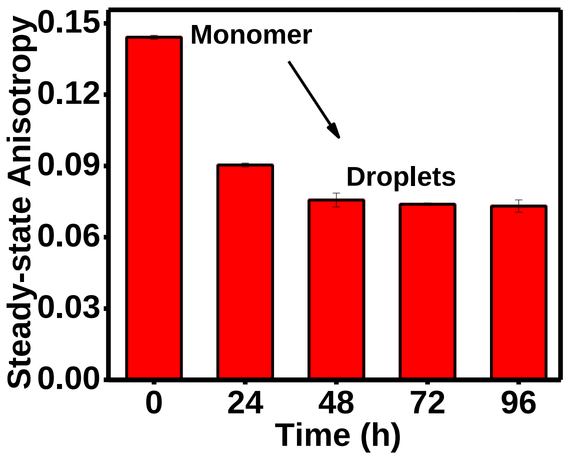


**Figure S5.** Steady-state fluorescence anisotropy of acrylodan-labeled tau K18 as a function of time during droplet formation.

**Table S1.** Values of the recovered Stern–Volmer constants ($K_{SV}$) obtained from fitting the plots shown in Figure 3A using Equation 4 and the bimolecular quenching rate constants ($k_q$) obtained using a relationship given by Equation 5.

| **Sample** | $\tau_0$ **(ns)** | $K_{sv}$ **(M$^{-1}$)** | $k_q$ **(M$^{-1}$ s$^{-1}$)** |
|---|---|---|---|
| **Free Fluorescein** | 3.94 ± 0.01 | 8.39 ± 0.18 | (2.1 ± 0.04) x $10^9$ |
| **Tau K18 Monomer** | 4.01 ± 0.03 | 4.24 ± 0.04 | (1.1 ± 0.01) x $10^9$ |
| **Tau K18 Droplet** | 3.91 ± 0.01 | 6.99 ± 0.10 | (1.8 ± 0.03) x $10^9$ |

**Table S2.** The typical parameters recovered from fitting (Equation 8) of the fluorescence anisotropy decay profiles shown in Figure 3C.

| **Sample** | $r_0$ | $\Phi_{fast}$ **(ns)** | $\beta_{fast}$ | $\Phi_{slow}$ **(ns)** | $\beta_{slow}$ |
|---|---|---|---|---|---|
| **Tau K18 monomer (0 h)** | 0.32 ± 0.002 | 0.46 ± 0.01 | 0.52 ± 0.004 | 3.18 ± 0.04 | 0.48 ± 0.004 |
| **Tau K18 Droplets (48 h)** | 0.34 ± 0.008 | 0.25 ± 0.01 | 0.72 ± 0.005 | 1.16 ± 0.01 | 0.28 ± 0.005 |
| **Tau K18 Droplets (72 h)** | 0.34 ± 0.03 | 0.24 ± 0.01 | 0.87 ± 0.003 | 1.13 ± 0.02 | 0.13 ± 0.007 |
